# Three distinct gamma oscillatory networks within cortical columns in macaque monkeys’ area V1

**DOI:** 10.1101/2023.09.30.560308

**Authors:** Eric Drebitz, Lukas-Paul Rausch, Esperanza Domingo Gil, Andreas K. Kreiter

## Abstract

A fundamental property of the neocortex is its columnar organization in many species. Generally, neurons of the same column share stimulus preferences and have strong anatomical connections across layers. These features suggest that neurons within a column operate as one unified network. Other features, like the different patterns of input and output connections of neurons located in separate layers and systematic differences in feature tuning, hint at a more segregated and possibly flexible functional organization of neurons within a column. To distinguish between these views of columnar processing, we conducted laminar recordings in macaques’ area V1 while they performed a demanding attention task. We found three separate regions with strong gamma oscillatory current source density (CSD) signals, one each in the supragranular, granular, and infragranular laminar domains. Their characteristics differed significantly in terms of their dominant gamma frequency and attention-dependent modulation of their gramma power and gamma frequency. In line, spiking activity in the supragranular, infragranular, and upper part of the granular domain exhibited strong phase coherence with their domain’s CSD signals but showed much weaker coherence with the other domains’ CSD signals. These results indicate that columnar processing involves a certain degree of independence between neurons in the three laminar domains, consistent with the assumption of multiple, separate intracolumnar ensembles. Such a functional organization offers various possibilities for dynamic network configuration, indicating that neurons in a column are not restricted to operate as one unified network. Thus, the findings open interesting new possibilities for future concepts and investigations on flexible, dynamic cortical ensemble formation and selective information processing.

## Introduction

The primate visual cortex consists of a complex network of neurons organized into multiple distinct areas that make up the visual processing pathways. Structurally, the cortex comprises six vertically stratified layers. Horizontally, it is organized into an array of vertical columns that include neurons of all six layers (Barbas et al., 2022; Mountcastle, 1997; Rockland and Ichinohe, 2004). Multiple lines of evidence suggest a functional role for these columnar circuits. Notably, neurons within the same column share similar preferences for specific stimulus features (Alonso, 2016; Kaschube et al., 2010). In the early visual cortex, neurons within a column have largely overlapping spatial receptive fields (RF) (Hubel and Wiesel, 1963, 1962; Mountcastle, 1957). Furthermore, in areas V1 and V2, neurons within a column share similar preferences for orientation, color, luminance, and ocular dominance (Garg et al., 2019; Lennie et al., 1990; Livingstone and Hubel, 1984; Peterhans and Von der Heydt, 1993; Poggio et al., 1975; Schiller et al., 1976).

This shared feature selectivity among neurons within the same column has been consistently observed across various areas of the visual cortex (Ahmed et al., 2012; Blasdel and Campbell, 2001; Franken and Reynolds, 2021; Gattass et al., 2005; Li et al., 2013; Niebur and Koch, 1994; Tootell et al., 1993; Vanduffel et al., 2002; Westerberg et al., 2021). Anatomically, the different types of neurons in the cortical layers exhibit characteristic patterns of strong vertical connections within columns, which have been proposed to implement a canonical microcircuit (Douglas and Martin, 2004). Based on these and related findings, the cortical column is widely recognized as a cohesive, functionally integrated local neuronal ensemble that operates as a single, unified entity (Capone et al., 2016; Douglas et al., 1989; Douglas and Martin, 2007; Hosoya, 2019; Jones and Rakic, 2010; Mountcastle, 1957).

However, despite important neuroanatomical and neurophysiological properties suggesting a unified mode of operation within a cortical column, further evidence suggests a potential for more segregated modes of operation. In such a scenario, neurons in different layers or groups of layers may not always operate in a fully integrated manner across the entire column but contribute to separate ensembles within a column. In line, neurons located in different layers of the same column can differ systematically in their feature tuning, for example, for orientation (Martinez et al., 2002; Ringach et al., 2002; Wang et al., 2020), stimulus size (Bijanzadeh et al., 2018), and other features (DiCarlo and Johnson, 2000; Hirsch et al., 2002; Hirsch and Martinez, 2006). The assumption of distinct functional roles for neurons in the column’s different layers is also supported by their different pattern of input and output connections (Harris and Mrsic-Flogel, 2013; Rockland, 2019) and theoretical considerations on their function, e.g., in grouping and attention (Grossberg, 2001; Hirsch and Martinez, 2006; Shushruth et al., 2012; Yazdanbakhsh and Grossberg, 2004). Thus, the question of whether neurons within a cortical column operate as a single, unified ensemble or in multiple, more segregated ensembles is open. Furthermore, in the latter case, the question would arise whether the configuration and interactions of such intra-columnar ensembles are fixed or might change on psychophysical timescales, possibly depending on the processing demands imposed by varying stimuli or behavioral tasks.

At the larger anatomical scale of cortical areas, experimental (Bosman et al., 2012; Drebitz et al., 2018; Gregoriou et al., 2012; Grothe et al., 2012) and theoretical studies (Aertsen et al., 1989; Borgers and Kopell, 2008; Briggs et al., 2013; Harnack et al., 2015; Segev and Rall, 1998; Tiesinga and Sejnowski, 2010) indicate that selective synchronization of oscillatory neuronal activity within the γ-band (30 – 100 Hz) provides a mechanism to configure neuronal ensembles dynamically and flexibly according to changing computational requirements. Changing γ-band synchronization between neurons thereby serves to modulate the functional connectivity within local neuronal populations (Aertsen et al., 1989; Azouz and Gray, 2003; Battaglia et al., 2012; Doesburg et al., 2007; Drebitz et al., 2018; Engel et al., 2013; Palmigiano et al., 2017) and allows for selective information routing and processing in neuronal systems despite of fixed anatomical connections (Fries, 2015; Kreiter, 2020).

Here, we apply this conceptual framework to the single cortical column to better understand its mode of operation and to improve the understanding of the internal dynamics of columnar processing and the information flow across columns. Assuming that a column operates as a single, unified entity, the concept predicts that neurons throughout the column synchronize to the same γ-rhythm and show uniform properties,regarding, for example, the γ-peak frequency, attention-dependent modulations of this frequency, or attention-dependent modulation of the oscillation’s amplitude. In contrast, if neurons in a column operate (at least temporarily) in functionally distinct ensembles, the neurons of each local ensemble should contribute to their ensemble’s temporal dynamics but not to a column-wide shared γ-rhythm. This ensemble-specific γ-oscillatory activity could differ in spectral characteristics and functionally relevant properties from those located in different columnar regions.

We tested these predictions using linear multi-contact probes to record neuronal activity simultaneously across the layers of cortical columns in area V1 of two macaque monkeys (*Macaca mulatta*). The animals performed a demanding attention task that required tracking the continuously changing shape of a previously cued stimulus. Subsequently, we computed the time-resolved current source density (CSD) to characterize the oscillatory γ-band activity (30-90 Hz) across layers.

We consistently found three distinct “hotspots” with strong γ-oscillatory activity during stimulus processing. They were separated from each other, with one located in the granular, one in the supragranular, and one in the infragranular layers of the column. The γ-oscillatory CSD signals at these three hotspots synchronized strongly with the spiking activity across the laminar domain in which they resided but less so with the spiking activity of other domains. Comparing the γ-band oscillations’ properties between the three regions showed marked differences. The dominant γ-frequencies and the strength of attention-dependent modulations of the γ-power differed significantly between the γ-oscillations at the three locations. Thus, our results indicate the presence of three different γ-oscillatory networks in the three major laminar domains of V1 columns, supporting the view that neurons within a cortical column can flexibly operate in multiple, at least partially segregated, ensembles.

## Results

To investigate the laminar pattern and characteristics of γ-oscillatory activity within the cortical column in area V1, we recorded neuronal activity using linear multi-contact probes with 100 μm inter-electrode spacing inserted perpendicular to the cortical surface. Concurrently, monkeys performed a demanding shape-tracking task while fixating a central fixation point (Fig. 1A/B). Before each trial, a spatial cue indicated the location of the behaviorally relevant stimulus. Animals started fixation and initiated the trial by pressing a lever. After a baseline period, up to four isoeccentric stimuli appeared simultaneously. Following a brief static presentation, all stimuli started to morph continuously from their initial shape into other complex shapes (Fig. 1B). Monkeys had to release the lever when, in the sequence of the cued stimulus, the initial shape reappeared. The reappearance could occur within two to four morph cycles, each lasting a second (Fig. 1B). Since one of the stimuli was located within the receptive fields of the neurons recorded within a V1 column, they processed either the behaviorally relevant target of attention or a non-relevant stimulus.

**Figure 1:**
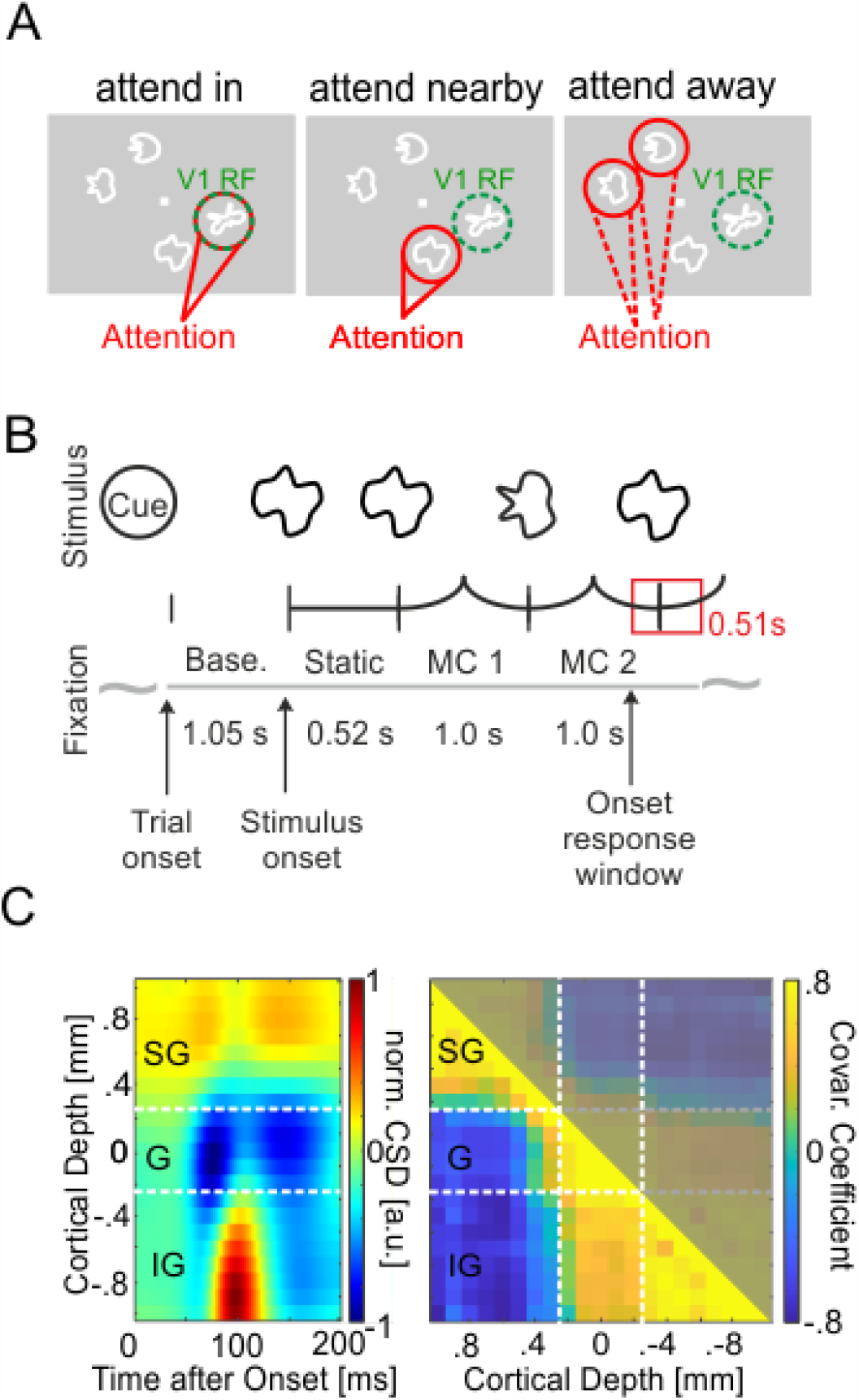
Task and layer discrimination. **A** Stimulus configuration of the three task conditions relevant for this study. Stimuli were shown at four locations on screen, one of them in the receptive fields (RFs) of the neurons recorded within the V1 column (indicated by a green dashed circle, not present on screen). Attention was cued by a spatial cue prior to the stimulus onset to one of the four locations (indicated in red). Attention could be focused on the recorded V1 site (attend in), the distractor close to the recorded V1 site (attend nearby), and at one of the two stimuli in the opposite hemifield (attend away). **B** Morphing sequence of a cued target stimulus with two morphing cycles (MCs). After cue presentation, animals initiated the trial by pressing a lever and holding fixation to the central fixation point (FP). Then, the cue disappeared, and after 1050 ms with only the FP shown (Baseline period, Base.), all stimuli were presented on screen. After a static presentation for 520 ms they started to continuously morph into other complex shapes within 1000 ms (MC). The reappearance of the initial shape at the cued stimulus location had to be signaled by the animals by releasing the lever within a response window of 510 ms (550 ms monkey I). **C** Estimation of the laminar position of electrodes. Left panel: Laminar profile of the CSD calculated from the average LFP in V1 in response to stimulus onset for one exemplary session of monkey I. White dashed lines mark the borders between the three major laminar domains: supragranular (SG), granular (G), and infragranular (IG) layers. Right panel: Average Pearson correlation between the CSD signals below 15 Hz during the baseline period of all combinations of two electrodes. Data taken from the same example session as in the left panel. Large and similar correlation values occurred between pairs of electrodes located within the same laminar domain.

### Three distinct hotspots of γ-band activity within V1 columns

To examine the vertical profile of oscillatory neuronal activity in the γ-band within V1 columns, we calculated the time-resolved CSD along a linear multi-contact probe and its power spectrum during MCs 2 and 3. The depth profiles of these power spectra for the individual sessions of both monkeys indicate the presence of several well-separated hotspots of high power in the γ-band along the recorded columns’ vertical axis (see Fig. 1A, left column for examples). The observations from individual sessions were confirmed after averaging the aligned and normalized power depth profiles of each monkey’s recording sessions. We observed a distinct peak within each of the three domains in 30 out of 32 recording sessions from both animals. However, in two sessions, either a discernible peak in the granular or infragranular domain was absent. On average, across all sessions, we found three distinct peaks of γ-power for each animal, one in each major laminar domain of the column, i.e., in the supragranular, the granular, and the infragranular layers. The laminar profile of the maximal γ-power at each electrode confirms the presence of these three peaks in both monkeys (Fig. 2B, right column, red curves). For comparison, we show the corresponding depth profiles of power spectra calculated from the LFP signals. Inspection of the profiles obtained for the individual sessions (see Fig. 2A, right column for examples) shows, on the one hand, that part of the peaks observed in the corresponding CSD profiles occur at similar locations and frequencies in the LFP profiles. On the other hand, the regions with enhanced γ-power are spatially much more extended than the more localized CSD peaks. In line, the average depth profile of LFP spectra (Fig. 2B, middle column) and the laminar profile of its maximal γ-power at each electrode (Fig. 2B, right column, blue curve) show enhanced γ-power across extended regions along the column, with little evidence for three separate hotspots.

**Figure 2:**
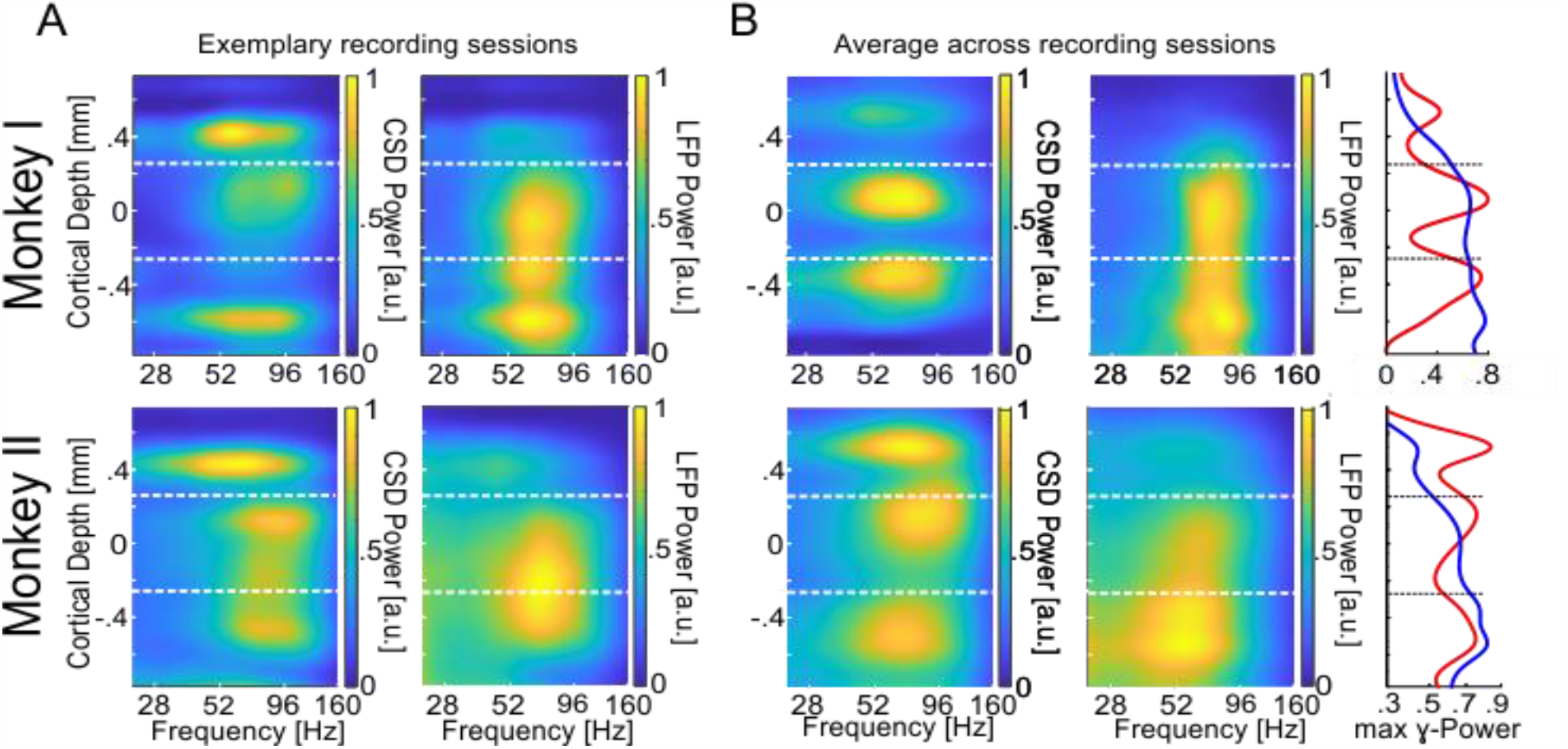
Cortical depth profile of CSD- and LFP-power in area V1. **A** The upper left panel displays the power (1/frequency corrected, maximum set to 1) of CSDs during all MCs 2/3 while attention was directed towards the rerecorded V1 RF (“attend in). The panel represents the average of an exemplary recording session of monkey I (same as in Fig. 1D). For the right panel, all conventions are equal, but power values are based on the LFP. The lower left panel displays the CSD power during the “attend in” condition for an exemplary recording site of monkey II. The lower right panel displays the same but for the LFP power. All other conventions are the same as for Monkey I. **B** The upper left and middle panels show the same (CSD left, LFP right) as in A (upper panels) for the average across all recording sessions. The lower left and middle panels display the same for Monkey II. The rightmost panels show the average γ-power of all recording sites within the γ-band. The red graphs represent the maxima of γ-CSD power, and the blue graphs the maxima of γ-LFP power. The vertical dashed lines (white and black) indicate the border between supragranular and granular layers (upper) and granular and infragranular layers (lower), respectively.

### γ-Frequency differs between γ-power hotspots in V1 columns

After identifying the three hotspots of γ-oscillatory activity in V1 columns, we asked whether they reflect three local groups of neurons operating during stimulus processing as a single, unified entity or, to some extent, functionally distinct ensembles. Therefore, we compared characteristics of neuronal γ-rhythmic activity between the three hotspots of V1 columns that could provide information on the mode of operation of the columnar networks. The precise frequency of the γ-oscillatory activity is one indicator that characterizes the dynamics of oscillatory networks. Identical γ-frequencies among the three hotspots’ CSDs would be expected if they all reflect a shared population rhythm across the entire column, while separate local γ-oscillatory networks are less likely to have identical γ-frequencies. We computed the time-resolved CSD signal for all trials in a session. Subsequently, we calculated the dominant γ-frequency for each of the three attentional conditions and each of the three hotspots of each session of both monkeys, using trials with four stimuli shown (see methods).

The results (Fig. 3A, left panel) show that the γ-frequencies at the three different hotspots differed significantly (Kruskal-Wallis test, *X*^2^(2) = 19.63, *p* = 5.47116*10^-5^, n_sup_= 78, n_gra_= 75, n_inf_= 75). After multi-comparison correction, the observed differences between the γ-frequency at the supragranular and granular hotspots and between the granular and the infragranular hotspots were significant (post hoc Dunn’s test, α < 0.05). The median γ-peak frequency of 70.7 Hz at the granular hotspot was about 4 Hz higher than at the supragranular hotspot with 67.0 Hz and about 5 Hz higher than at the infragranular hotspot with 65.3 Hz (Wilcoxon rank sum tests, *Z* = 2.958, *p* = 0.0031 and *Z* = 4.1534, *p* = 3.2758*10^-5^, respectively). No significant difference was found between the γ-frequency at the supragranular and infragranular hotspot.

**Figure 3:**
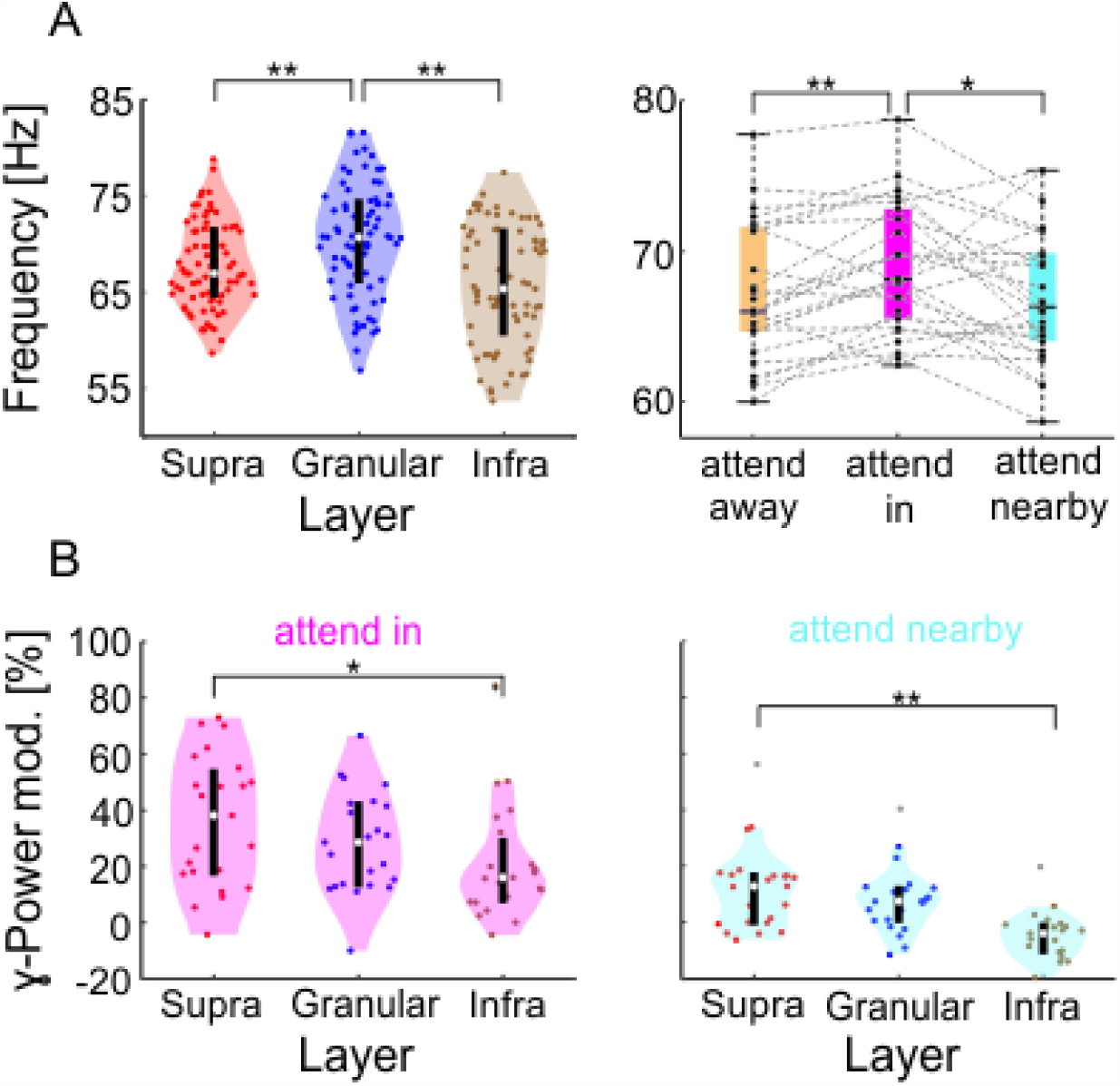
Comparison of spectral characteristics of the CSDs γ-oscillations at the hotspot of the supragranular-, granular, and infra-granular domain of both animals in V1. **A** The left panel shows the distributions of dominant γ-frequency across recording sessions. Highlighted in red, blue, and brown are the distributions for the supragranular, granular, and infragranular domains. The right panel shows the distribution of the dominant γ-frequency of the supragranular CSD signals for the three attentional conditions. **B** Distribution of the change in γ-power for the “attend in” (left) and “attend nearby” (right) conditions compared to the “attend away” condition. ** indicate significance at p < 0.01, and * at p < 0.05.

### Effects of attention differ between γ-power hotspots in V1 columns

If neurons within V1 columns operate to some extent in functionally distinct ensembles, they might be differentially affected by the direction of selective attention. Therefore, we investigated whether γ-frequency and γ-power differed between the three different attentional conditions (“attend in”, “attend nearby”, and “attend away”, see Fig 1 for illustration). We again used the trial-based, time-resolved CSD signal. However, now we separately analyzed the three attentional conditions in each session of both animals and determined the dominant γ-frequency for each hotspot and attentional condition. We found an effect of attention for the supragranular hotspot (Friedman test, *X*^2^(2) = 11.62, *p* = 0.003, n = 26). Attending the stimulus in the RF results in a median γ-frequency of 68.2 Hz, which is significantly higher than the 66.3 Hz when attending the closely neighboring stimulus or 66.0 Hz when attending stimuli in the opposite hemifield (Wilcoxon signed-rank tests, *Z* = 2.4255, *p* = 0.0459 and *Z* = 2.9589, *p* = 0.0093, respectively; p-values are Tukey-Kramer corrected). In contrast, no significant effect of attention on γ-frequency was observed for the granular and infragranular hotspots. Thus, the results indicate that attention differently affects the frequency of the γ-oscillatory activity at the three hotspots. Its major effect is a small frequency increase of about 2 Hz at the supragranular hotspot when attention is directed to the stimulus within the RF. In addition to the different effects of attention on γ-frequency at the three hotspots, their γ-power was also affected differently by attention. We again used the electrode centered at each hotspot. The relative increase of γ-power when attending the stimulus within the RF compared to stimuli in the opposite hemifield differed significantly between the three hotspots (Fig. 3B, left panel; Kruskal-Wallis test, *X*^2^(2) = 6.1191, *p* = 0.0469, n = 23). In this condition, the median attention-dependent increase of γ-power was 38.1% at the supragranular hotspot, 28.5% at the granular hotspot, and 16.0% at the infragranular hotspot. The difference between the attentional modulations at the supragranular and infragranular hotspots was statistically significant (post hoc Dunn’s test, α < 0.05 and Wilcoxon rank sum test, *Z* = 2.2848, *p* = 0.0223).

Similarly, directing attention to a stimulus near the RF as compared to the opposite hemifield also had differential effects on the γ-power of the three hotspots (Fig. 3B right panel; Kruskal-Wallis test, *X*^2^(2) = 22.8595, *p* = 1.0867*10^-5^). In this condition, the relative changes as compared to the “attend away” condition were generally smaller, with 12.8% at the supragranular hotspot, 7.5% at the granular hotspot, and -4.1% at the infragranular hotspot. Again, the difference between the attentional modulations at the supragranular and infragranular hotspots was statistically significant (post hoc Dunn’s test, α < 0.05 and Wilcoxon rank sum test, *Z* = 3.698, *p* = 2.6245*10^-4^).

Taken together, the three distinct hotspots of strong γ-oscillatory activity in cortical columns in V1 exhibit different characteristics not only in terms of their dominant γ-frequency but also in terms of the strength by which attention modulates their γ-frequency as well as their γ-power. This suggests that separate networks within these three major laminar domains engage in distinct γ-oscillations, which differ in frequency and the effect of attention on their spectral properties.

### Intra-columnar synchronization indicates separate γ-oscillatory networks

If V1 columns comprise three distinct networks, each exhibiting its own γ-oscillatory dynamics, it is expected that the activity of neurons that belong to the network would show stronger coupling to its γ-rhythm compared to the γ-rhythms of the other two networks. If, on the other hand, neurons across the entire column function as a unified ensemble with a single common γ-rhythm, then the activity of neurons throughout the column should exhibit similar coupling to this shared rhythm, regardless of where it is measured.

To test these predictions, we calculated the phase coherence (PhC) for each electrode site’s multi-unit activity with the three population rhythms. Subsequently, we investigated the columnar PhC patterns. As a measure of multi-unit activity, we used the entire spiking activity (ESA), which is a continuous signal and more sensitive to low amplitude spikes than measures requiring thresholding (Ahmadi et al., 2021a, 2021b; Drebitz et al., 2020, 2019). The ESA signals vary considerably in strength across cortical depth but are similar in spectral composition (see Fig. 4C for an exemplary ESA-power depth profile).

**Figure 4:**
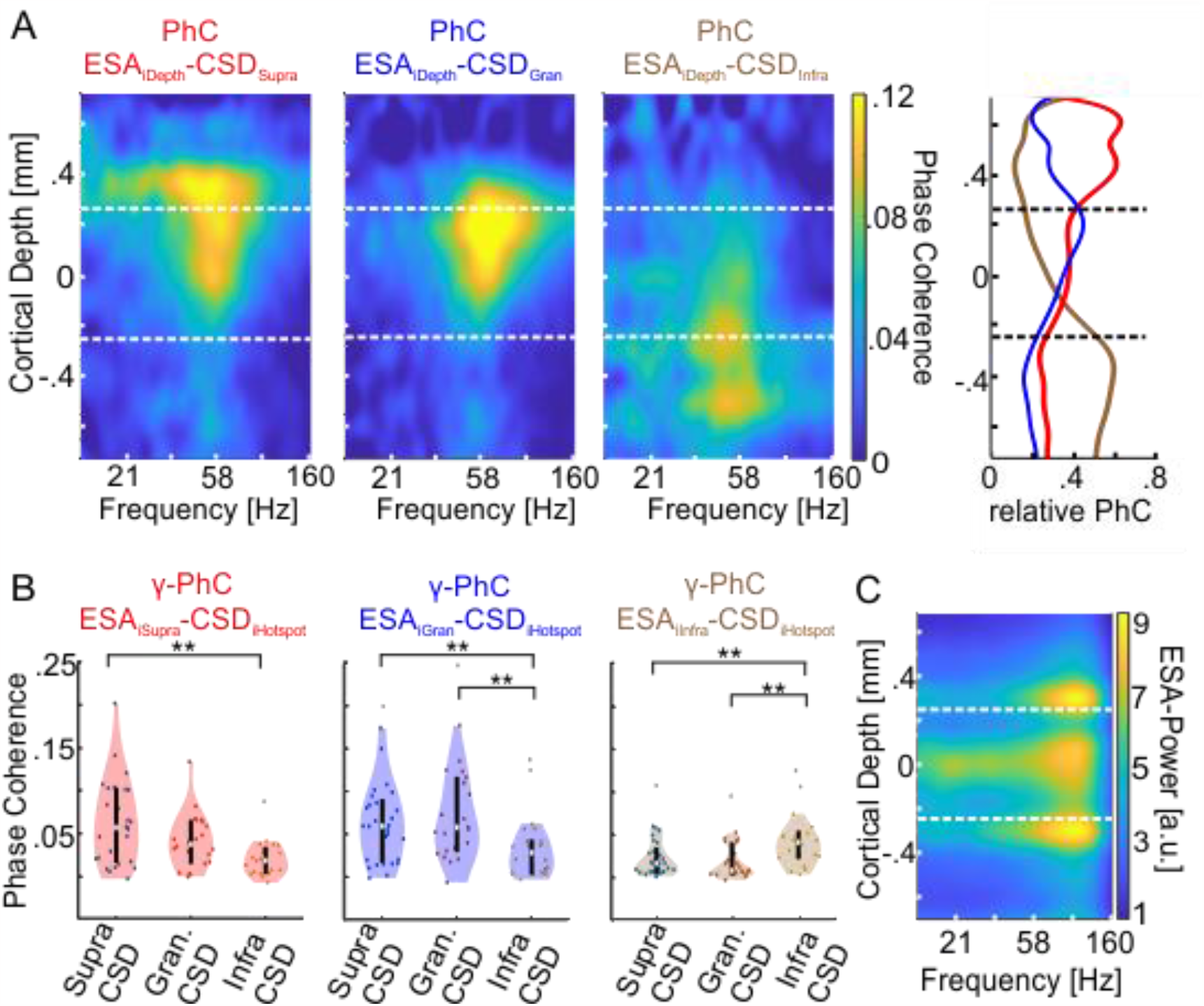
Depth profiles of phase coherence (PhC) of supragranular, granular and infra granular CSD with the entire spiking activity (ESA) at each electrode of both animals. **A** The left heatmap shows the average PhC-depth profile between the CSD signal of the γ-power hotspot in the supragranular domain with the ESA across all electrodes of all layers. The middle and right most heatmaps show the same for the CSD signals of the granular (blue title) and infragranular (brown title) hotspot, respectively. The vertical dashed lines indicate the borders between supragranular and granular layers and between granular and infragranular layers. The right most panel shows the average γ-PhC (28 – 78 Hz) across depth for each of the of the three CSD signals with ESA across depth, as ratio between each PhC value and the sum of the PhC values at the same depth. The lines are colored in the same hue as the corresponding heatmap’s title. **B** The left panel shows the distributions of the sessions’ average γ-PhCs between ESA the CSD signal of one of the three hotspots (depicted on the abscissa) across the supragranular electrodes. The white dot depicts the median γ-PhCs and the upper and lower edges of the black vertical line the 25^th^ and 75^th^ percentile, respectively. The middle and right panel show the same but for the respective combinations of granular and infragranular ESA signals with the CSD of the three hotspots’ γ-oscillatory activity. **C** The heatmap depicts the average ESA-power across a column for all “attend in”-conditions of an exemplary recording session of monkey II. * indicates significance between 0.01 < p < 0.05, ** indicates significance at p < 0.01.

The spectral patterns of PhC across cortical depth reveal that the CSD signal of each hotspot was maximally coherent with ESA from the same laminar domain with a peak in the γ-band (Fig. 4A, left three plots). Consequently, the set of ESA signals of a major laminar domain showed maximum γ-PhC with the CSD signal of the same domain and weaker γ-PhC with the CSD signals of the other two domains (Fig. 4B). All but two of these differences were statistically significant (Table 1).

**Table 1:**
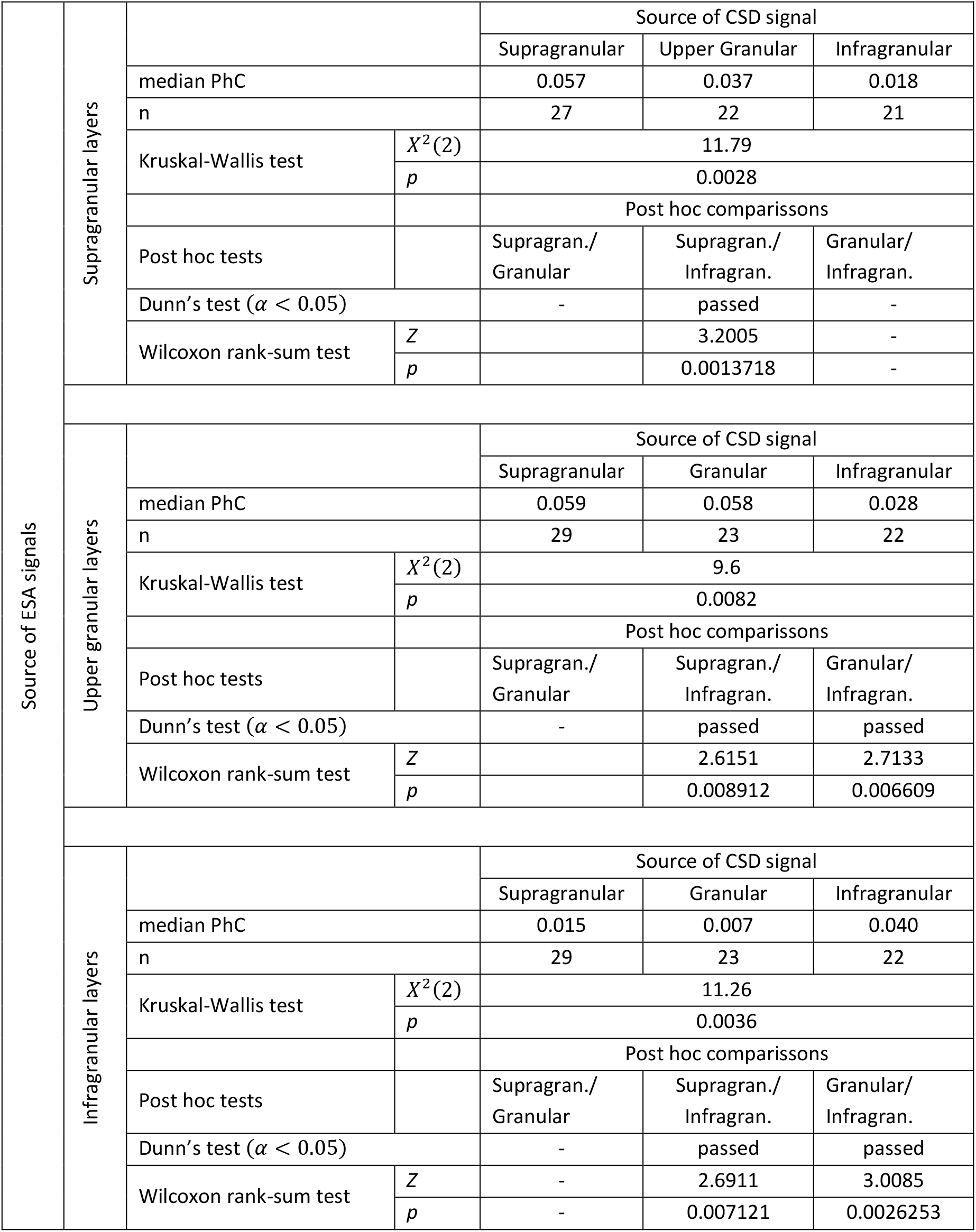
Statistical comparison of the strength of γ-PhC between the major laminar domain’s sets of ESA signals and the three CSD signals from γ-power hotspots.

Since ESA and its power in the γ-band varied strongly with cortical depth, the depth profile of PhC is correspondingly biased. Therefore, we calculated at each depth the mean PhC between ESA and each of the three CSD signals and averaged values within the γ-band. For each recording site, we normalized each of these three PhC-values by dividing with the sum of the three values (Fig. 4A, right panel).

The result reveals that ESA signals throughout the supragranular domain synchronize mostly with the γ-rhythm of the supragranular hotspot and in much smaller proportions with the γ-rhythm of the other two hotspots (Fig. 4A, right panel, red curve). At the border to the granular domain, the ESA’s PhC with the supragranular hotspot steps down to a lower level, which becomes even lower for ESA in the infragranular domain. A similar pattern occurs for the γ-rhythm at the infragranular hotspot. ESA signals throughout the infragranular domain synchronize mostly in similarly high proportions with this γ-rhythm (brown curve) and in much smaller proportions with the γ-rhythms of the other two hotspots. Above the infragranular domain’s border to the granular domain, the PhC of ESA with the infragranular γ-rhythm quickly declines. The fairly similar proportions of PhC by which the spiking activity of neurons throughout both laminar domains follow predominantly the γ-rhythm measured at the same domain’s hotspot suggests that the neurons across each domain belong to a unified neuronal ensemble and follow a domain-specific, common γ-rhythm.

The depth profile of ESA’s relative PhC with the γ-rhythm of the granular hotspot has a maximum close to the upper border of the granular domain, from where it gradually declines to the lower border and stays smaller throughout the infragranular domain (Fig. 4A, right panel, blue curve). Above the upper border, ESA’s relative PhC declines quickly. Despite the highest proportions of the PhC with the γ-rhythm of the granular hotspot resulting from spiking across the granular layer, this proportion is throughout the granular domain very similar to the proportion of PhC with the supragranular hotspot’s γ-rhythm. These patterns indicate that the γ-oscillatory network underlying hotspot of γ-power in the upper part of the granular domain might not extend homogeneously throughout the entire granular domain but is mainly located in its upper part.

In summary, the pattern of the PhC between the spiking activity across the column and the three hotspots of γ-power indicates three separate networks in the major laminar domains that engage in γ-oscillatory activity. Typically, the spiking activity of such a network’s neurons locks predominantly to their own network’s rhythm and less to the rhythms of the other networks.

## Discussion

Our study shows that V1 columns contain three distinct regions characterized by robust γ-oscillatory activity. These regions are localized in the supragranular, granular, and infragranular layers. The γ-oscillatory activities at these three locations exhibit notable differences in their characteristics. This concerns the dominant γ-frequency, which is significantly higher at the granular layer’s hotspot than at the supragranular and infragranular layers’ hotspots. Furthermore, attention increased the γ-frequency at the hotspot in the granular layers, while we observed no significant differences at the other two hotspots. Attention also changed the γ-power differentially across the three sites, with the strongest increase observed in supragranular layers and much smaller increases observed in infragranular layers. We also found that the spiking activity in the three laminar domains was more coherent with the CSD signal of the hotspot in the same domain but much less with the CSD signals of other domains.

Observing three distinct hotspots of high γ-power in CSD signals along the depth profile of V1 columns suggests that γ-oscillatory activity originates in three separate groups of neurons within a column. In good agreement, several previous studies showed a non-uniform distribution of γ-oscillatory activity in LFP across the cortical layers in area V1 (Han et al., 2021; Kienitz et al., 2021; Lowet et al., 2017; Maier et al., 2010; Spaak et al., 2012; Van Kerkoerle et al., 2014; Xing et al., 2012) and in extrastriate visual areas (Buffalo et al., 2011; Gieselmann and Thiele, 2022; Scheeringa and Fries, 2019) of macaques and in the human cortex (Csercsa et al., 2010). However, these LFP-based studies reported less than three peaks along the cortical depth and typically showed smoother spatial patterns that vary considerably between studies. Comparing LFP- and CSD-based depth profiles in the present study reveals that the limited spatial resolution inherent in LFP signals hinders the identification of all three γ-oscillators within a cortical column.

The presence of three hotspots of γ-oscillatory activity in the CSD raises the question of whether the three corresponding groups of neurons operate in a functionally fully integrated mode in one ensemble, forming one γ-oscillatory network throughout the column or whether they instead reflect three separate, even though flexibly interacting ensembles, each creating a γ-oscillatory network on its own. Several properties of the γ-oscillatory activity originating at the three hotspots suggest the latter possibility.

First, the dominant γ-frequency at the granular hotspot is significantly higher than the γ-frequencies of the supra- and infragranular hotspots. The corresponding neurons should follow the same rhythm if they belong to the same coherently oscillating ensemble. Even if this rhythm varies in instantaneous frequency, the average frequency should have been the same throughout the ensemble. Instead, the observed differences in γ-frequency suggest that the neurons from which the CSD signals at the three hotspots originate belong to different networks, each with its γ-oscillatory dynamics.

A second important difference between the neurons of these three networks concerns the effects of attention. Attending the stimulus processed by neurons throughout the column enhanced the γ-frequency of the granular network significantly but not of the two other γ-oscillatory networks. This finding may also help to explain discrepancies between previous studies in V1, which in part observed attention-dependent frequency changes in the γ-band of comparable effect sizes in V1 (Bosman et al., 2012; Das and Ray, 2018), while others report no significant changes in after accounting for multi-comparisons (Ray and Maunsell, 2010). Differences in the frequencies of recordings with conventional microelectrodes in different layers and the spatial averaging of LFP signals might contribute to these divergent outcomes. Ferro et al. (2021) reported a significant γ-frequency increase in V1 when pooling γ-frequency estimates based on LFP recordings from linear electrode arrays across all layers, but only slight differences in spectral signatures of attention between layers.

Besides the dominant γ-frequency of the CSD signals, selective attention also modulated the γ-power differently in the three networks. The attention-dependent increase ranged from 16 % at the infragranular hotspot to 38.1 % at the supragranular hotspot. Interestingly, prior research reported either an inconsistent or negligible impact of attention on V1 γ-power (Bosman et al., 2012; Buffalo et al., 2011; Ferro et al., 2021), observed even a suppressive effect (Chalk et al., 2010; Herrero et al., 2013), or showed comparatively small increase od γ-power in macaques (Rohenkohl et al., 2018) and humans (Magazzini and Singh, 2018). The particularly strong effects in the present study might partly reflect our task’s very high cognitive demands: continuously tracking the dynamic transformations of the target stimulus’s shape until its initial shape reappears in the presence of a closely spaced distractor, and while the luminance of all stimuli changed randomly and quickly. In contrast, previous tasks required detecting changes in color or orientation of a grating stimulus or the sudden appearance of a bright spot on a darker bar. At the same time, the stronger effect might be due to the high spatial resolution of CSD signals, which reduces the diminishing of power modulations by spatial averaging with less modulated or unmodulated signals.

However, the pronounced differences observed for the attention-dependent modulation of γ-frequency and γ-power between the three hotspots of a V1 column fit the expectation for separate γ-oscillatory networks with distinct functional roles that are, therefore, separately and differently influenced by selective attention. Direct evidence for three networks, each generating its γ-rhythm, comes from the PhC analysis between the highly local spiking activity acquired throughout the entire column depth and each CSD signal from the three hotspots of γ-power. While the raw PhC between, e.g., the supragranular hotspot’s CSD signal and the ESA at different electrodes varies abruptly, strongly depending on the strength and spectral characteristics of the different ESA signals, the proportions between an ESA signal’s PhCs with the three different CSD signals reveal the spatial organization of the three γ-oscillatory networks. The comparatively high and similar proportion of PhC between spiking activity throughout supra- and infragranular layers with the corresponding hotspots in these laminar domains indicates that neurons throughout each domain engage in a local γ-rhythm. The same holds for the γ-rhythm of the granular hotspot that predominantly reflects the rhythm of neurons’ spiking activity in the granular layers’ upper part. The γ-oscillatory CSD signals at the three hotspots reflect these population rhythms. They are specific for the three different networks, as indicated by the differences in their γ-frequency and the weaker PhC between neurons’ spiking activity and the population rhythms of the other networks compared to their own network’s rhythm.

These results indicate a notable degree of independence between the dynamics of the γ-oscillatory networks, a trait commonly associated with weakly coupled γ-oscillatory networks (Hoppensteadt and Izhikevich, 1998; Kopell and Ermentrout, 2002; Palmigiano et al., 2017; Schuster and Wagner, 1990). Typically, such dynamics emerge in networks with stronger connections in local groups of neurons that engage in oscillatory dynamics and weaker connections between such local oscillatory networks (Kopell and Ermentrout, 1986; Palmigiano et al., 2017; Wang, 1995). The intracolumnar connectivity in V1 resembles such an architecture. Taking the supragranular layers as an example, most of their neurons’ synaptic inputs (∼55%) originate from intralaminar connections. In contrast, less than half of this percentage (∼20%) arises from granular layers, with a mere ∼10% originating from infragranular layers. The remaining input is traced back to other cortical areas (Binzegger et al., 2009; Dantzker and Callaway, 2000; Lindén et al., 2011; Potjans and Diesmann, 2014). Thus, the known connectivity of the intracolumnar microcircuitry shows critical properties expected if the γ-oscillatory networks in the different laminar domains behave like weakly coupled oscillatory networks.

It seems intriguing that local columnar networks within V1 exhibit weakly coupled oscillatory behavior even though they are thought to transmit and process stimulus information sequentially from granular to supragranular and infragranular layers without obvious necessity for selection and signal gating. However, this disposition toward weak coupling yields valuable degrees of independence that can enhance information processing efficiency:

1. A gradual modulation of phase coherence and phase difference between, e.g., the granular and the supragranular networks’ γ-oscillations could implement an adjustable gain factor for the input signal processed in a V1 column before it is passed to subsequent processing stages. Such adjustable gain factors are required, e.g., by divisive normalization models for flexibly weighting different input signals (Carandini and Heeger, 2012; Kreiter, 2020). Furthermore, such a flexible gain modulation within a cortical column may provide another possibility to modulate signal- and information routing at the columnar level and is supported by the observation of attention-dependent changes of Granger causality between columnar networks (Ferro et al., 2021).
2. The same basic mechanisms might serve to change the pattern of functional connectivity between the column’s laminar networks and, thereby, the proportions between the weights of intra-columnar inputs and extra-columnar inputs from neighboring columns as well as from bottom-up or top-down inputs originating outside of V1. For example, reduced coupling between granular and supragranular layers could allow for stronger weighting of the supragranular network’s interactions with neighboring columns via horizontal connections. This could promote the local binding of stimulus features for integrating stimulus information from the classical RF and its surround (Lowet et al., 2017; Vinck and Bosman, 2016).
3. It is known that transmitting signals along a chain of weakly coupled oscillators enhances robustness to perturbations and decreases the likelihood of cascade failures (Cosp et al., 2004; Csaba and Porod, 2020; Rode et al., 2019). The intra-columnar signal pathways across γ-oscillatory networks in different layers could also benefit from this effect.

Future research will be required to understand the possible functional role of the flexible modulation of functional connectivity between different layers in columnar signal- and information processing in the context of different demands due to changing stimulus constellations and task requirements.

We identified three distinct, local γ-oscillatory networks within V1 columns, each in a different laminar domain. Their γ-oscillatory activity revealed notable differences concerning the modulation of their spectral features by selective attention. The phase coherence patterns within and between the networks show characteristics of weakly coupled oscillatory networks. This allows their functional connectivity to change by changing coherence and phase relations between their γ-oscillations. It implies various possible mechanisms that could support cortical information processing at the columnar level. The findings indicate that neurons within a V1 column do not operate as a single, unified ensemble but in multiple γ-oscillatory ensembles with considerable potential for flexible reconfiguration of the columnar functional networks. Because of the anatomical and histological similarities across the cortex, we anticipate that these findings will likely extend to other cortical areas.

## Methods

### Materials and Methods

#### Surgical procedures

Two adult male macaque monkeys (Macaca mulatta) were implanted with titanium head posts and recording chambers above areas V1/V2 under aseptic conditions. All procedures were approved by the local authorities (Senator für Gesundheit, Bremen, Germany) and followed the German Animal Welfare Act (TierSchG) and the European Council directive 2010/63/EU for laboratory animal care and use.

#### Data acquisition and experimental setup

During recording sessions, monkeys sat in a custom-made primate chair in front of a 20-inch CRT monitor (monkey I: 93 cm, monkey II: 95 cm) with a display resolution of 1280 x 1024 pixels and a refresh rate of 100 Hz (ELSA Ecomo 750). Eye position was monitored by video-oculography (IScan Inc., Woburn, MA, USA). See Drebitz et al., 2018, for details of the experimental setup.

Semi-chronic recordings in cortical area V1 were performed with linear multi-contact probes. Each probe had 16 or 32 electrodes with a center-to-center spacing of 100 μm and a pointy tip (ATLAS Neuroengineering bvba, Leuven, Belgium). The iridium oxide-coated electrodes had an impedance of 0.25 MΩ at 1 kHz and a diameter of 15*15 μm. A small custom-made microdrive was fixed to a grid in the recording chamber to hold and move the probe in a direction perpendicular to the cortical surface. After each recording session, the probe was retracted until the tip was located just below or in the dura mater. It was repositioned within the cortex at the start of each subsequent session. We conducted up to five sessions before removing the microdrive from the recording chamber, replacing the linear probe if necessary, and placing it in a new position in the chamber. At the initial probe insertion, neuronal signals were monitored to detect the penetration of the dura and the entering of the individual electrodes into the cortex to control for accurate depth placement. Using an automatic visual stimulation and data analysis procedure, we simultaneously mapped the receptive fields (RF) of all recording sites along the probe based on their induced ESA and the induced γ-band power in the LFP in response to a moving bar (see Drebitz et al., 2019 for details). Given the semi-chronic recording approach, the receptive field (RF) locations remained consistent across successive sessions for which the same probe was used at the same place. Nevertheless, prior to the recordings in these successive sessions, we confirmed the previously observed RF locations by manually mapping the RFs using a bar stimulus.

Neuronal signals were amplified 20,000-fold for monkey I and 16,000-fold for monkey II with a wideband preamplifier (MPA32I) and a programmable gain amplifier (PGA 64, 1 – 5,000 Hz; Preamplifier and PGA, Multi Channel Systems GmbH, Reutlingen, Germany) and subsequently digitized with a 12-bit ADC at 25 kHz sampling rate. The signals of all recording sites were referenced against the titanium recording chamber (27 mm diameter) that was in contact with bone and intracranial tissue.

#### Visual task paradigm

During V1 recordings, the animals performed a demanding shape-tracking task requiring the animals to attend to the continuously changing shape of a target stimulus and respond to the reappearance of its initial shape while fixating a central fixation point (FP) (Drebitz et al., 2018) in the presence of distractor stimuli. The appearance of this shape at one of the distractor stimuli had to be ignored. The stimulus configuration for the task encompassed up to four stimuli, isoeccentrically arranged around the FP (Fig. 1A). One of the four stimuli was centered on the overlapping RFs of the neurons recorded in the V1 column. A second stimulus resided close to but outside of the V1 RFs. The other two stimuli were placed at point mirrored locations in the opposite visual hemifield with respect to the central FP (Fig. 1A). For estimating neuronal responses without a nearby stimulus and to control that the nearby stimulus is located outside the RFs, in half of the trials either the stimulus in the RF or the stimulus nearby were not shown. Before the trial onset, a spatial cue was presented in the form of an annulus with a diameter of 1°, a line width of 0.25°, and a luminance of 3.8 cd/m^2^. This cue indicated the specific location where the behaviorally relevant stimulus would appear during the subsequent trial. The two stimulus locations within or nearby the recorded V1 RFs were cued with 30 % probability, respectively. The two stimulus locations in the opposite hemifield were cued with 20 % probability, resulting in three task conditions. Attention is focused either on the stimulus within the RFs of the recorded V1 site (“attend in” condition), on the stimulus close to the recorded V1 site (“attend nearby” condition), or at a stimulus in the opposite hemifield (“attend away” condition, see Fig. 1B).

The cue appeared with the FP (square, 0.15°x0.15°, 2.45 Cd/m^2^) on the dark monitor (background luminance: 0.03 Cd/m^2^) and vanished after the monkeys initiated the trial by pressing a lever. After a baseline period of 1050 ms, the complex-shaped stimuli (∼1°-1.5° diameter, line width 0.25°) appeared, and following a static period (Fig. 1B), they started to change their shape, morphing continuously into other complex shapes. One morph cycle lasted one second, i.e., the time for morphing from one shape into the next. When the initial shape of the cued target stimulus the target shape reappeared at the end of MCs 2, 3, or 4, monkeys had to release the lever within a response window (Fig. 1B, red rectangle) of 550 ms (510 ms), starting 380 (310) ms before this MC ends and terminating 170 (200) ms after its end for monkey I (monkey II). Shapes for each stimulus were drawn randomly from a set of 12 shapes for monkey I, two of which could become the target shape, and from a set of 8 shapes for monkey II, all of which could become the target shape. To heighten task difficulty, the stimulus luminance changed randomly with each frame of the screen. For this, a random, integer gray pixel value was selected before each frame update [0 255] corresponding to luminance changes from 0.02 – 45.4 Cd/m^2^, on average 3.8 Cd/m^2^ (for details: Grothe et al., 2018; Lisitsyn et al., 2020). From trial initiation until lever release, monkeys had to hold their gaze within a window of 1° diameter centered on the fixation square. If they did so and responded within the response window to the reappearance of the target shape at the cued stimulus location, they received a water reward. Otherwise, the trial ended without reward.

## Data Analysis

All analyses described below were performed with custom MATLAB scripts running on MATLAB v. 2020a (MathWorks, Natick, MA, USA). The violin plots were created using an open-source MATLAB script available at https://github.com/bastibe/Violinplot-Matlab (Bechtold, 2016).

### Estimation of Current Source Density (CSD)

To determine the time-resolved CSD signals for each trial, we first extracted the local field potentials (LFP) by filtering the recorded broadband neuronal signals with a low-pass FIR filter (pass: <160 Hz, stop: 300 Hz, suppression: 80 dB) in forward and backward direction. The resulting LFPs were subsequently downsampled to 1000 Hz and used for calculating the CSD signal with the spline inverse CSD method (iCSD), as described by Pettersen et al., 2006. This approach supposes that the recorded LFPs result from current sources within the cortical column that are evenly distributed in cylindrical discs stacked on top of each other and that the CSD varies smoothly in z-direction along the column’s center axis (perpendicular to the cortical layers) as described by a set of cubic polynomials. For the present analyses, we assumed a disc radius of 500 μm and an isotropic medium with a conductivity of 0.4 S/m (Logothetis et al., 2007). The depth profile of the CSD was calculated and spatially smoothed with a Gaussian filter (SD 200 μm, in a finite window of 1000 μm) for each point in time using the open-access software provided by and available at Pettersen, 2005. This provided for each electrode and trial a CSD signal as a function of time. For the discrimination of layers based on source and sink locations, we averaged the LFP within the first 250 ms starting after stimulus onset (see Fig. 1B) across the trials of all conditions with a stimulus in the RF before computing the CSD using the spline iCSD method.

### Estimation of Entire Spiking Activity (ESA)

To investigate the synchronization between spiking activity and the CSD signals, we estimated the entire spiking activity (ESA), which is a continuous signal and which does not reject spikes with small amplitude by thresholding (for a comparison of MUA and ESA, see Drebitz et al., 2019). We obtained the ESA by first filtering the neuronal data with a high-pass FIR filter (pass: >400 Hz, stop: 300 Hz, suppression: -80 dB) in forward and backward directions to obtain the high-frequency components of the signal. The high-passed signal was full-wave-rectified and low-pass filtered with the same FIR filter as used for obtaining the LFP in forward and backward directions. Subsequently, the ESA signal was downsampled to 1000 Hz.

### Spectral analysis

The spectral decomposition of CSD, LFP, and ESA signals was performed by convolving them with complex Morlet’s wavelets, 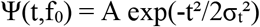 exp(2iπf_0_t), with σ_f_= 1/πσ_t_ and *f*_0_/*σ*_*f*_ = 6 (Kronland-Martinet et al., 1987; Taylor et al., 2005; Torrence and Compo, 1998). The wavelets were normalized such that the total energy was 1, requiring the normalization factor A to be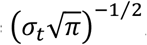. The central frequencies f_0_ of the daughter wavelets were set between 5 and 160 Hz based on a scheme of Torrence and Compo (1998). The time-frequency resolved power spectral density was then calculated by taking the square of the absolute value of the result of the convolution and dividing it by the Nyquist frequency of 500 Hz (Drebitz et al., 2018; Taylor et al., 2005). If not stated otherwise, we corrected for the 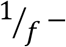 bias of the PSD by multiplying the PSD values with the central frequency of the corresponding wavelet. Time-frequency resolved power spectral density was calculated separately for each session and attentional condition. Power spectra were calculated by taking the mean of all power values for the same frequency across time within MCs 2 and 3.

### Estimation of the dominant γ-Frequency

The dominant γ-frequency for the CSD signal of the central recording site of a domain’s hotspot was calculated for each trial based on the average γ-oscillatory cycle period. Only γ-oscillations with more than average amplitude during MCs 2 and 3 were considered. We first filtered the CSD signals with a broad bandpass filter (FIR-filter, passband: 35-120 Hz, stop frequencies: 25 Hz and 140 Hz, suppression: -20 dB) in forward and backward directions. The band-limited CSD signals were Hilbert transformed to calculate instantaneous phase *φ*(*t*) and amplitude *A*(*t*) of the γ-oscillations (Le Van Quyen et al., 2001; Rosenblum et al., 2001). The time course of the amplitude *A*(*t*) was smoothed with a Gaussian filter (*σ* = 10 ms). In time spans of at least 40 ms duration in which the amplitude continuously exceeds the median amplitude value during MCs 2 and 3 of the trial, we determined the points in time when the γ-oscillations’ troughs occurred, i.e., when the instantaneous phase *φ*(*t*) crosses *π*. Since this happened usually between the phase values sampled at discrete times, we determined the crossing of *π* using linear interpolation. The period lengths of the γ-oscillatory cycles were calculated as the time difference between successive throughs. Few periods incompatible with frequencies within the filter’s pass band (i.e., >1/35 s or <1/120 s) were discarded. For each session and each attentional condition, we determined the median γ-cycle period based on all periods calculated as described above. We calculated the dominant frequency as the reciprocal of the median gamma cycle period.

### Estimation of Phase Coherence (PhC)

We calculated the phase coherence (PhC), which is also called phase locking value (PLV), by calculating the resultant vector of phase differences between ESA and CSD signals over *N* trials. The phase values *φ*_*CSD*_(*t*, *f*) and *φ*_*ESA*_(*t*, *f*) at times *t* and frequency *f* were derived from the convolution of CSD- and ESA-signals with complex Morlet’s wavelets (see Spectral Decomposition). The PhC for N trials is given by:

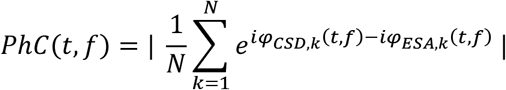

To allow for comparison of measurements with different numbers of trials, we corrected the PhC values for their trial-number-dependent bias by subtracting the expected value (*EV*) of the PhC for N trials as in the actual analysis, but with random phase relations, which is given by: 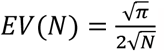 (Grothe et al., 2012).

### Layer Identification and electrode alignment

To identify the position of the individual electrodes of a probe with respect to the cortical layers and align the electrodes between recording sessions according to their cortical position, we combined two procedures: First, we used the spatiotemporal pattern of the CSD calculated with the spline iCSD method from the average field potential response to stimulus onset as described above and identified the border between layer III and layer IV based on the inversion of the CSD’s polarity from the source closest to the cortical surface to the initial sink below (see Figure 1D, left panel for an example). Second, we calculated a correlation matrix containing for each pair of electrodes the average Pearson correlation between their CSD signals during the baseline period (Fig. 1B). For this purpose, in each trial, the time-resolved CSD signals of all electrodes were low-pass filtered with an FIR filterpass: < 15 Hz, stop: 20 Hz, suppression: 35 dB) in forward and backward directions within the time window from 100 ms after the start of the baseline period until 100 ms before its end. Subsequently, the Pearson correlation coefficient between these low-frequency CSD signals was calculated for each trial and pair of electrodes. Finally, the median value of each electrode pair’s correlation coefficients across trials provided an entry into the matrix of median correlation coefficients (see Fig 1D, right panel for an example). The correlation matrix usually delivered a sharp decrease in correlation around the border of layers III/IV and helped to refine the results of the first procedure. Recording sites above the border between layers III and IV were categorized as supragranular. Recording sites located within 0.5 mm below this border were categorized as granular, and sites further below as infragranular (Ferro et al., 2021; Maier et al., 2010; Van Kerkoerle et al., 2014).

### Data selection

We recorded neuronal data in 14 sessions for monkey I and 18 sessions for monkey II. If not stated otherwise, data from MCs 2 and 3 of correctly performed trials was analyzed. If an MC terminated with the reappearance of the initial shape of the target stimulus, only its data until 150 ms before the animal released the lever was used.

For the analysis of the PSD of CSD and LFP data across the cortical depth, we used all available sessions for each monkey.

Subsequent analyses required the signals from the center of the CSD’s γ-band-power hotspot within each of the three major domains of the column, i.e., the supragranular, the granular, and the infragranular layers. For this purpose, we first computed in all 32 sessions the 1/f corrected (see Methods: Spectral analysis) power spectrum of the CSD in MCs 2 and 3 for each of the five electrode positions in the granular layer, the five adjacent electrode positions above in supragranular layers, and the five electrode positions below in infragranular layers across all correctly performed trials on the attend-in condition with four stimuli present. Then, we computed the mean spectrum across sessions for each of these 15 electrode positions of each monkey. Taking the mean spectrum across these spectra for the 15 electrode positions allowed defining the borders of the frequency window for γ-power calculation as the width of the PSD peak in the γ-band at half height of its maximum value (38.4 Hz to 118 Hz in both animals). Finally, we computed the mean γ-power within these borders from each electrode’s power spectrum in each session. In the resulting depth profile of CSD γ-power of each recording session, we searched each of the column’s three domains for a discrete peak flanked at both sides by electrode positions of the same domain with lower γ-power. In the few sections without such a discrete peak, we searched for a peak hidden in the flank of another larger peak in neighboring regions, as indicated by a flattening in such a flank, but without a local maximum. Therefore, we calculated the gradient of the γ-power between the section’s electrodes. In the case of a local minimum of the gradient at a rising flank or of a local maximum of the gradient in a falling flank, we selected from the electrode pair with the gradient’s extremum the first or second electrode, respectively. If both approaches failed, no electrode was selected for the investigated domain of the column and session.

For the analysis of phase coherence between CSD and ESA, we used in all sessions as the source for CSD signals the three (or fewer) electrodes that we located at the centers of the hotspots of CSD γ-band power in the three main sections of the recorded column (as described above). To avoid analyzing phase coherence based on recording sites with no or only weak responses to the stimulus, we excluded electrodes with a CSD γ-power during MCs 2 and 3 less than 2.5 times the CSD γ-power during the baseline period. As a source for ESA signals, we selected two-thirds of all electrodes in each session with the strongest ESA γ-power in MCs two and three for the “attend in” condition. For analyzing attention-dependent modulations of the power and dominant frequency in the γ-band, we used the 26 sessions with a behavioral performance of 65% or better (disregarding trials terminated by breaking fixation).

### Statistics

The statistical procedures to determine significance are denoted alongside each respective comparison in brackets. These procedures included Kruskal-Wallis tests for assessing significant differences between multiple independent sample populations (at p<0.05), initially conducted on unpaired data to address the multiple comparisons issue due to comparisons between three hotspots. Subsequently, Dunn’s tests were applied to identify significantly distinct sample groups (p<0.05). For the groups where Dunn’s test indicated significance, we conducted a one-sided post-hoc Wilcoxon rank-sum test, providing the pertinent statistical parameters.

For the statistical evaluation of paired data, we initially conducted a Friedman’s test to determine whether there were statistically significant differences among the related groups. Subsequently, we employed post-hoc Wilcoxon-signed rank tests (two-sided). To control for the type I errors due to multiple comparisons, we applied the Tukey-Kramer correction to adjust the p-values.

## Acknowledgments

This work was funded by the Deutsche Forschungsgemeinschaft (DFG, German Research Foundation) - 331514942.

In addition, we thank Dr. Iris Grothe for valuable discussions on the manuscript. We also thank Aleksandra Nadolski, Peter Bujotzek, and Katja Taylor for training and technical assistance. Furthermore, we want to acknowledge Katrin Thoß and Ramazani Hakizimana for their diligent animal care.

